# MAIT cell deficiency exacerbates neuroinflammation in P301S human tau transgenic mice

**DOI:** 10.1101/2025.01.03.631124

**Authors:** Yuanyue Zhang, Zhi Yang, Xiaosheng Tan, Na Jiang, Gaoyuan Cao, Qi Yang

**Affiliations:** Child Health Institute of New Jersey, Rutgers Robert Wood Johnson Medical School, New Brunswick, NJ, 08901, USA; Rutgers Institute for Translational Medicine and Science, Rutgers Robert Wood Johnson Medical School, New Brunswick, NJ, 08901, USA; Department of Pediatrics, Rutgers Robert Wood Johnson Medical School, New Brunswick, NJ, 08901, USA

**Keywords:** Neurodegeneration, Tau pathology, P301S mice, MAIT cells, innate-like T cells, mucosal associated invariant T cells

## Abstract

The role of immune cells in neurodegeneration remains incompletely understood. Our recent study revealed the presence of mucosal-associated invariant T (MAIT) cells in the meninges, where they express antioxidant molecules to maintain meningeal barrier integrity. Accumulation of misfolded tau proteins are a hallmark of neurodegenerative diseases. The role of MAIT cells in tau-related neuroinflammation and neurodegeneration, however, remains unclear. Here we report that the meninges of P301 mutant human tau transgenic mice had increased numbers of MAIT cells, which retained their expression of antioxidant molecules. *Mr1^−/−^*P301S mice that lacked MAIT cells exhibited increased tau pathology and hippocampus atrophy compared to control *Mr1*^+/+^ P301S mice. Adoptive transfer of MAIT cells reduced tau pathology and hippocampus atrophy in *Mr1^−/−^* P301S mice. Meningeal barrier integrity was compromised in *Mr/^−/−^*P301S mice, but not in control *Mr1*^+/+^ P301S mice. A distinctive microglia subset with proinflammatory gene expression profile (M-inflammatory) was enriched in the hippocampus of *Mr1^−/−^* P301S mice. The transcriptomes of the remaining microglia in these mice also shifted towards a proinflammatory state, with increased expression of inflammatory cytokines, chemokines, and genes related with ribosome biogenesis and immune responses to toxic substances. The transfer of MAIT cells restored meningeal barrier integrity and suppressed microglial inflammation in the *Mr1^−/−^* P301S mice. Together, our data indicate an important role for MAIT cells in regulating tau-pathology-related neuroinflammation and neurodegeneration.

## Background

Aberrant aggregation of misfolded microtubule-associated protein tau is a prominent feature of neurodegenerative disorders [1]. Previous studies have shown that the density of neurofibrillary tangles, which are intracellular aggregates of hyperphosphorylated tau, correlates with the severity of cognitive decline in Alzheimer’s disease patients [2, 3]. Recent research suggested a complex interplay between tau pathology and neuroinflammation. Notably, microglial inflammation has been observed in mice carrying human tau mutants associated with degenerative tau pathologies, suggesting that tau-induced pathologies can trigger neuroinflammation [4]. Conversely, reactive microglia and its products may also exacerbate tau pathology and associated neurodegeneration [5–8].

Moreover, recent studies suggest that non-microglial immune cells, such as lymphocytes, may play a role in modulating tau pathology-related neuroinflammation and neurodegeneration [5]. CD8^+^ T cells have been detected infiltrating the brain parenchyma in Alzheimer’s disease patients, as well as in 3D human neuroimmune culture derived from stem cells from AD patients and in mouse models with neurodegenerative tau pathology [5, 9–12]. Increased infiltration of CD8^+^ T cells into 3-D human neuroimmune culture led to increased microglial activation, neuroinflammation and neurodegeneration [9]. Depletion of CD8^+^ T cells has also been shown to reduce tau pathology and prevent tau-mediated neurodegeneration in mouse models [5]. However, the roles of other lymphocyte subsets in regulating tau pathology and neurodegeneration remain largely unexplored.

MAIT cells are the predominant type of innate-like T cells in humans [13]. The T cell receptor [14] of MAIT cells recognizes microbiota-derived vitamin B2 metabolites presented by MR1 (MHCI-related)-expressing antigen presenting cells (APC) [15–17]. Thus, the development of MAIT cells requires microbiota and MR1-expressing thymocytes for positive selection [15],[16, 17]. However, fully matured MAIT cells can survive and function in the absence of MR1/TCR signaling, highlighting innate-like property [15–17]. We recently found that MAIT cells were present in the leptomeninges and secreted many antioxidant proteins to protect meningeal barrier integrity [18]. MAIT cell deficient mice exhibited notable meningeal barrier leakage, which inflamed the brain and disrupted cognitive function [18]. These data together demonstrate an important role for MAIT cells in protecting meningeal barrier integrity, and suggest that activate maintenance of the meningeal barrier integrity by MAIT cells is essential for restricting neuroinflammation and for protecting cognitive function. MAIT cells may also regulate amyloid β clearance in mouse models [19]. However, whether MAIT cells play a role in regulating tau pathology-related neuroinflammation and neurodegeneration remains unknown.

In this study, we crossed MAIT cell-deficient mice (*Mr1*^−/−^) with P301S mutant human tau transgenic mice to investigate the potential roles of MAIT cells in regulating tau pathology, neuroinflammation, and neurodegeneration. We found that MAIT cells are present in the meninges of P301S mice and maintain high expression of antioxidant molecules. Our results indicate that the absence of MAIT cells leads to meningeal leakage, increased neuroinflammation, and exacerbated tau pathology and hippocampal atrophy in P301S transgenic mice. These findings suggest that MAIT cells play a role in modulating tau pathology and associated neuroinflammation and neurodegeneration.

## Methods

P301S mice [4] and control mice were obtained from JAX laboratory. *Mr1*^−/−^ mice were described previously [18, 20]. P301S mice were bred with *Mr1*^−/−^ mice at animal facility of Rutgers University. Age and sex matched female and male mice of 7-8 month-old were used in this study. All animal experiments were performed according to protocols approved by the Institutional Animal Care and Use Committee at Rutgers University.

### Flow cytometric analysis and fluorescence activated cell sorting (FACS)

Flow cytometric analysis and FACS sorting of meningeal MAIT cells were carried out as previously described [18]. Briefly, the meninges were carefully isolated from the outer dorsal cerebrum and inner calvaria surface. The tissue was incubated for 20 minutes at 37°C in HBSS containing 0.2 mg/mL Liberase TM [21] and 0.1 mg/mL DNase I [21] to facilitate digestion. The digested tissue was passed through a 70 µm cell strainer to generate a single-cell suspension. For flow cytometry, the cells were stained with MR1 tetramers along with antibodies targeting surface antigens for 30 minutes at 22°C. The mouse MR1 5-OP-RU tetramers were obtained from the NIH Tetramer Core Facility. The MR1 tetramers were produced by the NIH Tetramer Core, as authorized for distribution by the University of Melbourne [22]. To define and characterize MAIT cells, surface markers used in the analysis included anti-TCRβ (H57-597), anti-CD3ε (145-2C11), anti-B220 (RA3-6B2), anti-NK1.1 (PK136), anti-CD11b (M1/70), anti-CD45 (104), anti-IL18R (A17071D), anti-Thy1.2 (53-2.1), and anti-IL7R (A7R34). Anti-B220, anti-NK1.1, and anti-CD11b antibodies were used to exclude B cells, NK cells, and myeloid cells to enhance MAIT cell identification. These antibodies were obtained from Biolegend, Thermo Fisher, or BD Biosciences. In flow cytometry and FACS sorting for QPCR analysis, meningeal MAIT cells were identified as CD45^+^ CD11b^−^ B220^−^ NK1.1^−^ MR1tetramer^+^ CD3/TCRβ^+^. In some experiments, anti-IL18R, anti-Thy1.2, and anti-IL7R antibodies were used for further characterize MAIT cells. Dead cells were excluded using DAPI (Thermo). For adoptive transfer experiments, no anti-CD3 or anti-TCRβ antibodies were included to prevent cell activation, and donor MAIT cells were identified as CD45^+^ Thy1^hi^ IL-18R^hi^ MR1tetramer^+^.

For analysis and sorting of hippocampal microglia, hippocampal tissue was processed using the papain-based Pierce™ Primary Neuron Isolation Kit (Thermo), according to the manufacturer’s protocol. The cells were passed through a 70 µm cell strainer and stained with surface markers, including anti-CD11b (M1/70) and anti-CD45 (104), before being sorted by FACS. Microglia was identified as CD11b^med/low^CD45^med/low^ cells as we previously described [23–25]. Intracellular cytokine staining was performed as we previously described [20, 23–28]. Specifically, after digestion, cells were cultured with serum free RPMI medium for 4 hours, and 1 µM Monensin monensin was added for the last 2 hours. The cells were then stained for surface markers to identify microglia, followed by intracellular staining with anti-TNF antibody (MP6-XT22). Intracellular cytokine staining was performed using the Cytofix/Cytoperm Kit (BD) as per the manufacturer’s instructions.

Flow cytometry was performed using a 4-laser LSRII (BD Biosciences) or a 4-laser Cytek Aurora, and cell sorting was performed on a Cytek Aurora CS cell sorter.

### Examination of Tau pathology and hippocampus weight

To examine tau pathology, immunofluorescence staining was performed with AT8 antibodies, which recognize tau phosphorylation at the Ser202 and Thr205 positions [29, 30]. Mice were first perfused with 50 mL of PBS, followed by 50 mL of 4% paraformaldehyde. The brains were fixed overnight in 4% paraformaldehyde at 4°C and then placed in 30% sucrose at 4°C until the tissue sank. The brains were then embedded in OCT and stored at −80°C. The brain sections were cut at 40 µm using a CM1850 cryostat (Leica). The sections were incubated with AT8 antibodies (Thermo Fisher) at a dilution of 1:100 in 0.1% BSA overnight at 4°C, followed by labeling with Donkey anti-Mouse IgG (H+L) Highly Cross-Adsorbed Secondary Antibody, Alexa Fluor 594. Images of the slides were captured using a BZ-X800 all-in-one fluorescence microscope (Keyence) with the BZ-X800 software.

For measurement of the hippocampus weight, the hippocampal tissue was carefully isolated from each hemisphere, weighed in an EP tube, and the weight was calculated by subtracting the weight of the tube before and after adding the hippocampal tissue.

### Adoptive transfer of MAIT cells

For adoptive transfer, meningeal MAIT cells (CD45^+^Thy1^+^IL18R^hi^MR1-tetramer^+^) from 7-month-old P301S mice were sorted by FACS, and 1000 cells were transferred to 6 weeks old *Mr1*^−/−^ P301S mice intravenously. To minimize undesired cell activation, CD3 and TCR antibodies were not included for purification of donor cells for adoptive transfer. Centrifugation, FACS sorting and preparation for adoptive transfer, were carefully performed on ice or 4°C. Meningeal leakage assays, immunofluorescence assays and hippocampus weight measurement were performed when recipient mice were around 7-8 months old.

### Single nucleus RNA-seq

For single nucleus RNA-seq (snRNA-seq), the hippocampus was quickly isolated from fresh mouse brains on ice, and placed in Hibernate E medium (Gibco) supplemented with B27 (GibcO) and 1% GlutaMAX (Gibco). The tissue was homogenized using a Dounce grinder in pre-cooled RNase-free lysis buffer (10 mM Tris-HCl, 10 mM NaCl, 3 mM MgCl2, 0.1% NonidetTM P40). The homogenization was performed with 15 strokes using a loose pestle, followed by 7 strokes using a tight pestle. The tissue was then lysed on ice for 15 minutes, and the solution was filtered through a 30 µm cell strainer. The nuclei were washed, and myelin was removed using myelin removal kit (Miltenyi) according to the manufacturer’s instructions. Libraries were prepared using 3’ GEM gene expression kit (10xgenomics) following the manufacturer’s instructions. Libraries were sequenced using a Novaseq. The scRNA-seq data were analyzed using the R package Seurat 4.3.0. Normalization of the data was achieved using the SCTransform function within Seurat. The R package DoubletFinder 2.0.3 was used to remove the doublets. Cell clustering was performed using Uniform Manifold Approximation and Projection (UMAP). Microglia/macrophage population was sorted in silicon based on expression of positive markers (e.g. *Itgam, Ptprc*) and the absence of markers associated with other immune cells (e.g. T/B/NK/ILC cells, monocytes, granulocytes). The microglia/macrophage population was then separated into distinct microglia subsets using UMAP. Gene markers for each microglia subset and differentially expressed genes between microglia subsets of *Mr1*^−/−^ P301S and control *Mr1*^+/+^ P301S mice were identified using a threshold of false discovery rate of <0.05. The Gene Set Enrichment Analysis (GSEA) algorithm was utilized to access the enriched gene sets from differential genes. In brief, the differential gene expression of targeted groups was calculated with the R package limma 3.56.2. Then the differential genes were preranked by log2FoldChange and subjected to the R package clusterProfiler 4.8.3. Results with p value<0.05, NES>1 and False discovery rate<0.25 were considered significant for GSEA analysis.

### Multiplex cytokine assays and QPCR analysis

Multiplex cytokine assays were performed as we previous described [23, 25]. The brain was homogenized with 1ml of PBS containing protease inhibitors using a tissue homogenizer. Concentrations of proinflammatory cytokines in brain homogenates were measured using LegendPlex kits, a kit for bead-based multiplex assays (Biolegend) [31], following the manufacturer’s instructions.

For QPCR analysis, microglia were sorted by FACS from the hippocampus. mRNA was extracted using the RNeasy Plus Mini Kit (Qiagen), following the manufacturer’s protocols. Complementary DNA (cDNA) was synthesized using SuperScript II Reverse Transcriptase (Thermo). qPCR was carried out using ABI pre-optimized TaqMan probes (thermos).

### Statistical analysis

For snRNA-seq data, Wilcoxon rank-sum test was used to determine significance of differentially expressed genes. False discovery rate less than 0.05 was considered significant. A Mann-Whitney U test was used to compare the differences in the percentages of subsets in each mouse for snRNA-seq data, and a p-value of less than 0.05 was considered significant. For GSEA analysis, results with p value<0.05, NES>1 and False discovery rate<0.25 were considered significant. For other experiments, student T test, or ANOVA test with Tukey’s post-hoc test was used to determine the difference between two groups. A p-value of less than 0.05 was considered significant.

## Results

### MAIT cells are present in the meninges of P301 mutant human tau transgenic mice and retain expression of anti-oxidant molecules

Our recent work indicates that MAIT cells are present in adult mouse meninges where they secrete anti-oxidant molecules to protect meningeal barrier integrity. We performed flow cytometry analysis to examine MAIT cells in the meninges and brain parenchyma regions of P301 mutant human tau transgenic mice to determine whether tau pathology might affect MAIT cells abundance and distribution. MAIT cells were detected in the meninges of 7-month-old P301S mice and their numbers were increased compared to age-matched control wildtype mice (Fig. 1A, 1B). MAIT cells did not further infiltrate into the brain parenchymal regions such as hippocampus and cortex of P301S mice (Fig.1B). Meningeal MAIT cells in 7-month old wildtype mice expressed MAIT-characteristic markers such as IL7R, IL18R, and CD3ε, verifying their identity (Fig.1C). They expressed comparable mRNA levels of anti-oxidant molecules such as *Selenop*, *Selenof, Fth1, Tmsb4x*, and *Psap*, as MAIT cells in wildtype mice, indicating that MAIT cells in P301S mice might retain anti-oxidant activities (Fig. 1D). Thus, MAIT cells are increased in the meninges of P301S mice and retain expression of anti-oxidant molecules.

**Figure 1:**
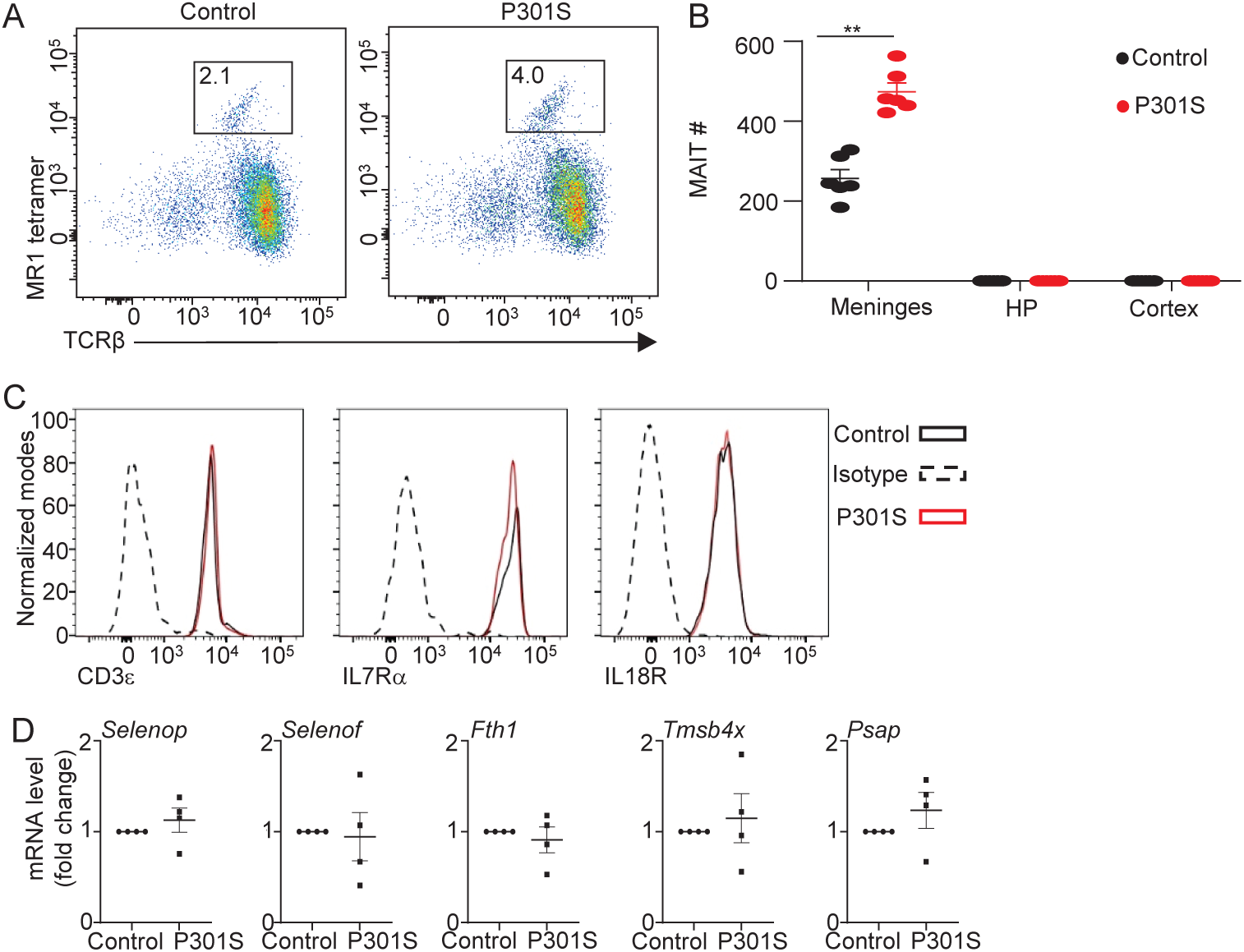
MAIT cells are present in meninges of P301S mice and retain expression of antioxidant molecules. (**A**) Representative flow cytometry profiles of MAIT cells in the meninges of 7-month old P301S mice and control wildtype mice. Plots were pre-gated on CD45^+^CD11b^−^B220^−^Thy1^+^ cells. (**B**) Numbers of MAIT cells in the meninges, hippocampus (HP), and cortex of 7-month old P301S mice and wildtype mice. (**C**) Representative flow cytometry profiles showing expression of the indicated genes by meningeal MAIT cells in P301S mice. Plots were pre-gated on meningeal MAIT cells in P301S mice. (**D**) mRNA levels of the indicated genes in meningeal MAIT cells of 7-month old P301S mice and control wildtype mice. Data are from 6 mice per group pooled from 2 independent experiments (B), or 4 independent experiments with 5 mice per group pooled experiments (D). Dat were normalized to *Gapdh*, and values in the control groups were set as 1.

### MAIT cell deficiency exacerbates tau pathology and hippocampus atrophy in P301S mice

Because MAIT cell development requires Mr1-expressing thymocytes for positive selection, *Mr1*^−/−^ mice lack MAIT cells due to failure in early development. To examine potential role of MAIT cells in regulating tau pathology, we crossed *Mr1^−/−^* mice, which lacked MAIT cells, with P301S mice. As expected, MAIT cells were absent in the meninges of *Mr1^−/−^* P301S mice (Fig. 2A, 2B). We examined tau pathology by immunofluorescence staining with AT8 antibody, which detects tau phosphorylation at the Ser202 and Thr205 positions [29, 30]. In mice bred in our animal facility, *Mr1^−/−^*P301S mice exhibited significantly increased p-tau pathology compared to control *Mr1*^+/+^P301S mice as early as 7-month-old (Fig. 2C, 2D). At 8 months old, *Mr1^−/−^* P301S mice showed a drastic increase in p-tau pathology compared with control *Mr1*^+/+^P301S mice (Fig. 2C, 2D). Adoptive transfer of MAIT cells partially restored MAIT cell numbers in the meninges (Fig. 2E), and repressed p-tau pathology in *Mr1^−/−^* P301S mice (Fig. 2F). Together, these data indicate that MAIT cell deficiency exacerbates tau pathology.

**Figure 2:**
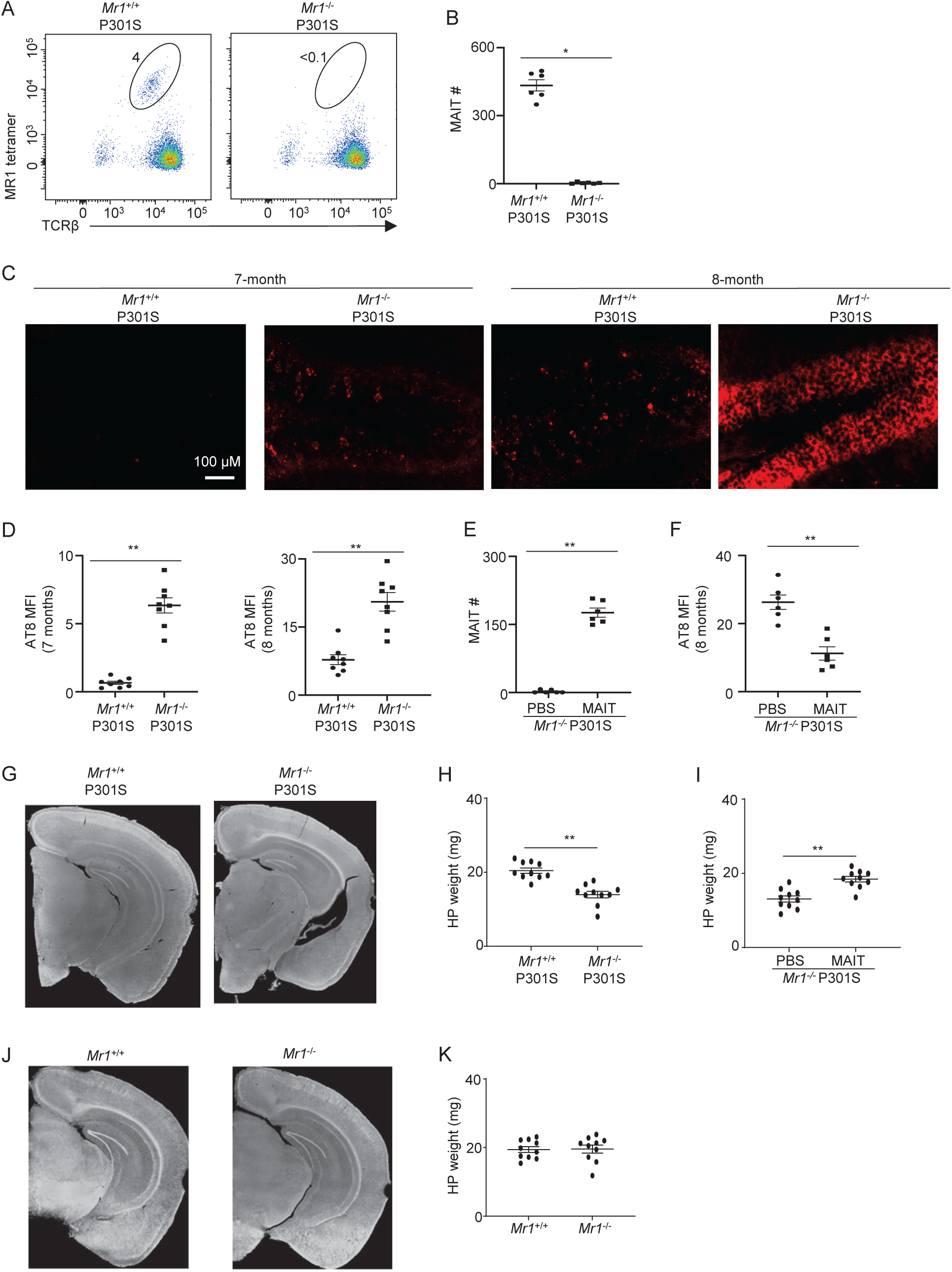
MAIT cells repress tau pathology and neurodegeneration in P301S mice. (**A**) Representative flow cytometry profiles of MAIT cells in the meninges of 7-month old *Mr1*^−/−^ P301S mice and control *Mr1*^+/+^ P301S mice. Plots were pre-gated on CD45^+^CD11b^−^B220^−^Thy1^+^ cells. (**B**) Numbers of MAIT cells in the meninges of 7-month old *Mr1*^−/−^ P301S mice and control *Mr1*^+/+^ P301S mice. (**C**) Immunofluorescence staining of p-Tau (antibody clone AT8) in 7-month and 8-month *Mr1*^−/−^ P301S mice and control *Mr1*^+/+^ P301S mice. Shown is hippocampus DG region. (**D**) Mean fluorescence intensity of p-Tau in the hippocampus of 7-month and 8-month *Mr1*^−/−^ P301S mice and control *Mr1*^+/+^ P301S mice. (**E**) MAIT cells were transferred to 6-week-old *Mr1*^−/−^ P301S mice. Number of MAIT cells in meninges of recipient mice was measured by flow cytometry analysis, when the mice were 8-months old. (**F**) Mean fluorescence intensity of p-Tau in the hippocampus of 8-month *Mr1*^−/−^ P301S mice with adoptive transfer of MAIT cells or PBS. (**G**) Representative images of hippocampus of 7.5-month *Mr1*^−/−^ P301S mice and control *Mr1*^+/+^ P301S mice. (**H**) Weight of the hippocampus in 7.5-month *Mr1*^−/−^ P301S mice and control *Mr1*^+/+^ P301S mice. (**I**) Weight or the hippocampus in 7.5-month *Mr1*^−/−^ P301S mice with adoptive transfer of MIT cells or PBS. (**J**) Representative images of hippocampus of 7.5-month *Mr1*^−/−^ P301S mice and control *Mr1*^+/+^ P301S mice. (**K**) Weight of the hippocampus in 7.5-month *Mr1*^−/−^ and control *Mr1*^+/+^ mice. Data are from 6-10 mice per group, 2 independent experiments.

We next examined the role of MAIT cells in regulating tau-pathology-associated neurodegeneration. At 7.5 months of age, most control *Mr1*^+/+^P301S mice did not yet exhibit notable hippocampus atrophy in mice bred in our animal facility (Fig. 2G, 2H). In contrast, *Mr1^−/−^* P301S mice displayed significant hippocampus atrophy at this age, indicating that the absence of MAIT cells may exacerbate tau-related neurodegeneration (Fig. 2G, 2H). The transfer of MAIT cells alleviated hippocampus atrophy in *Mr1^−/−^* P301S mice, verifying that MAIT cells repress neurodegeneration in P301S mice (Fig. 2I). Of note, *Mr1^−/−^* mice without human tau mutant transgene did not show hippocampus atrophy, indicating that the increased neurodegeneration in *Mr1^−/−^* P301S mice was associated with tau pathology (Fig. 2J, 2K). Together, these data suggest that MAIT cells restrict tau-pathology-associated neurodegeneration.

### MAIT cell deficiency leads to meningeal leakage and increased microglial inflammation in P301S mice

We aim to understand the mechanisms by which MAIT cells suppress tau pathology and neurodegeneration in P301S mice. Previous studies indicate that increased microglial inflammation can worsen tau pathology and neurodegeneration in P301S mice [5, 32–34]. Our previous work suggests that MAIT cells protect meningeal barrier integrity via secretion of antioxidant molecules, which could help prevent noxious substances from enteringthe brain parenchyma and triggering enhanced microglial inflammation. We thus examined meningeal barrier integrity and microglial activities in *Mr1^−/−^* P301S mice. At 7-months of age, a remarkable leakage in the meningeal barrier was observed in 7-month old *Mr1^−/−^*P301S mice, but not in control *Mr1*^+/+^ P301S mice (Fig. 3A, 3B). Transfer of MAIT cells restored meningeal integrity in *Mr1^−/−^* P301S mice, indicating that MAIT cells help prevent meningeal leakage in P301S mice (Fig. 3C). Notably, flow cytometry analysis revealed that microglia in *Mr1^−/−^* P301S mice expressed markedly high levels of the proinflammatory cytokine TNF (Fig. 3D-3F). Transfer of MAIT cells reduced TNF expression from microglia in *Mr1^−/−^*P301S mice, suggesting that MAIT cells supress neuroinflammation in P301S mice (Fig. 3G). These results indicate that the absence of MAIT cells leads to meningeal leakage and increased microglia inflammation in *Mr1^−/−^* P301S mice.

**Figure 3:**
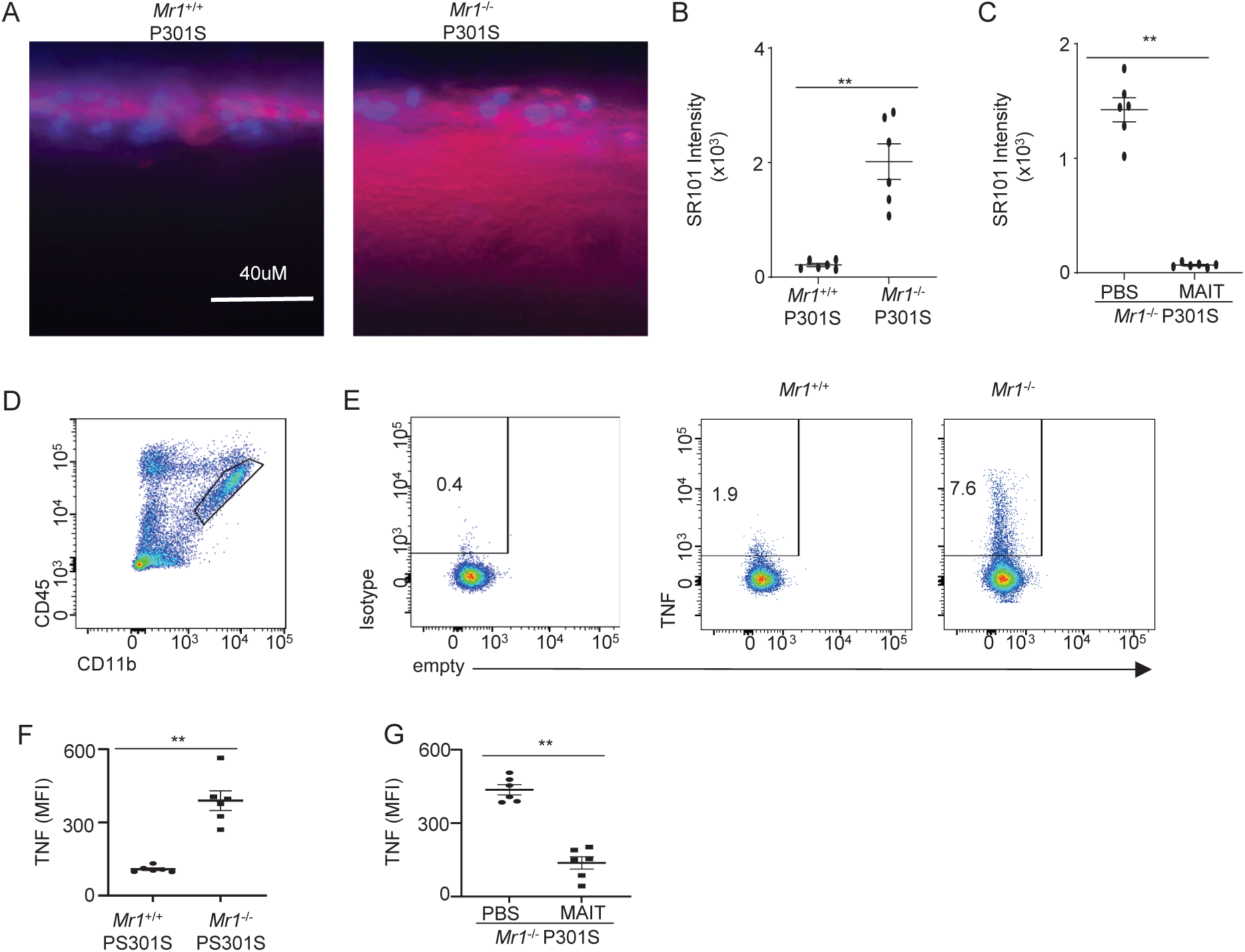
MAIT cells protect meningeal barrier integrity and repress proinflammatory cytokine expression from microglia. (**A**) Brain vibratome sections were obtained from 7-month-old *Mr1*^−/−^ P301S mice and control *Mr1*^+/+^ P301S mice with transcranial administration of SR101. Representative imaging of sections at 300µM to 600µM lateral to bregma. (**B**) Quantification of fluorescence intensity of SR101 at 50µM below the leptomeningeal cells in the brain vibratome sections of 7-month-old *Mr1*^−/−^ P301S mice and control *Mr1*^+/+^ P301S mice. (**C**) MAIT cells were transferred to 6-week-old *Mr1*^−/−^ P301S mice. Coronal brain vibratome sections were obtained in mice with transcranial administration of SR101, when mice were 7-month-old. Quantification of fluorescence intensity of SR101 at 50µM below the leptomeningeal cells in mice with or without adoptive transfer of MAIT cells. (**D**) Representative flow cytometry profile showing gating strategy of microglia in the hippocampus. (**E**) Representative flow cytometry profiles showing expression of TNF in microglia in the hippocampus of 7-month-old *Mr1*^−/−^ P301S mice and control *Mr1*^+/+^ P301S mice. (**F**) Mean fluorescence intensity of TNF in microglia in the hippocampus of 7-month-old *Mr1*^−/−^ P301S mice and control *Mr1*^+/+^ P301S mice. (**G**) Mean fluorescence intensity of TNF in the hippocampus of 7-month *Mr1*^−/−^ P301S mice with adoptive transfer of MAIT cells or PBS. Data are from 6 mice per group, representative of 2 independent experiments.

### Characterization of the transcriptomes of microglia in *Mr1^−/−^* P301S mice

We next performed single-nucleus RNA-seq to examine the transcriptomes of microglia in the hippocampus of *Mr1^−/−^* P301S mice and control *Mr1*^+/+^P301S mice. Unsupervised UMAP divided hippocampal microglia/macrophage cells into several subsets (Fig. 4A). The three most abundant microglia subsets differed in their expression of ribosome biogenesis genes, and were thus termed “M_ribo_hi”, “M_ribo_med”, and “M_ribo_lo” (Fig. 4A, 4B). In addition, a few minor subsets were detected (Fig. 4A). M_proliferating expressed high levels of proliferation markets such as *Ki67* (Fig. 4B). M_ISG expressed high levels of interferon stimulated genes (ISG) such as *Ifi44, Oasl2, Ifit2, Ifi204,* and *Ifit3*. M_magi2 and M_nrg3 were named by their expression of characteristic genes Magi2 and Nrg3 (Fig. 4B). Notably, a distinct subset of microglia appeared, expressing a high level of proinflammatory cytokines such as *Tnf*, *Il1a*, and *Il1b*, as well as proinflammatory chemokines such as Ccl3 and Ccl4 (Fig. 4A, 4B). We named this subset Inflammatory Microglia (M_inflammatory) (Fig. 4A, 4B). *Mr1^−/−^* P301S mice had significantly higher percentages of M_inflammatory subset, compared to control *Mr1*^+/+^ P301S mice (Fig. 4C, 4D). The other microglia/macrophage subsets did not show significant difference in percentages between *Mr1^−/−^* P301S mice and control *Mr1*^+/+^ P301S mice (Fig. 4C, 4D). Thus, *Mr1^−/−^* P301S mice had accumulation of a distinctive subset of microglia expressing high levels of proinflammatory genes.

**Figure 4:**
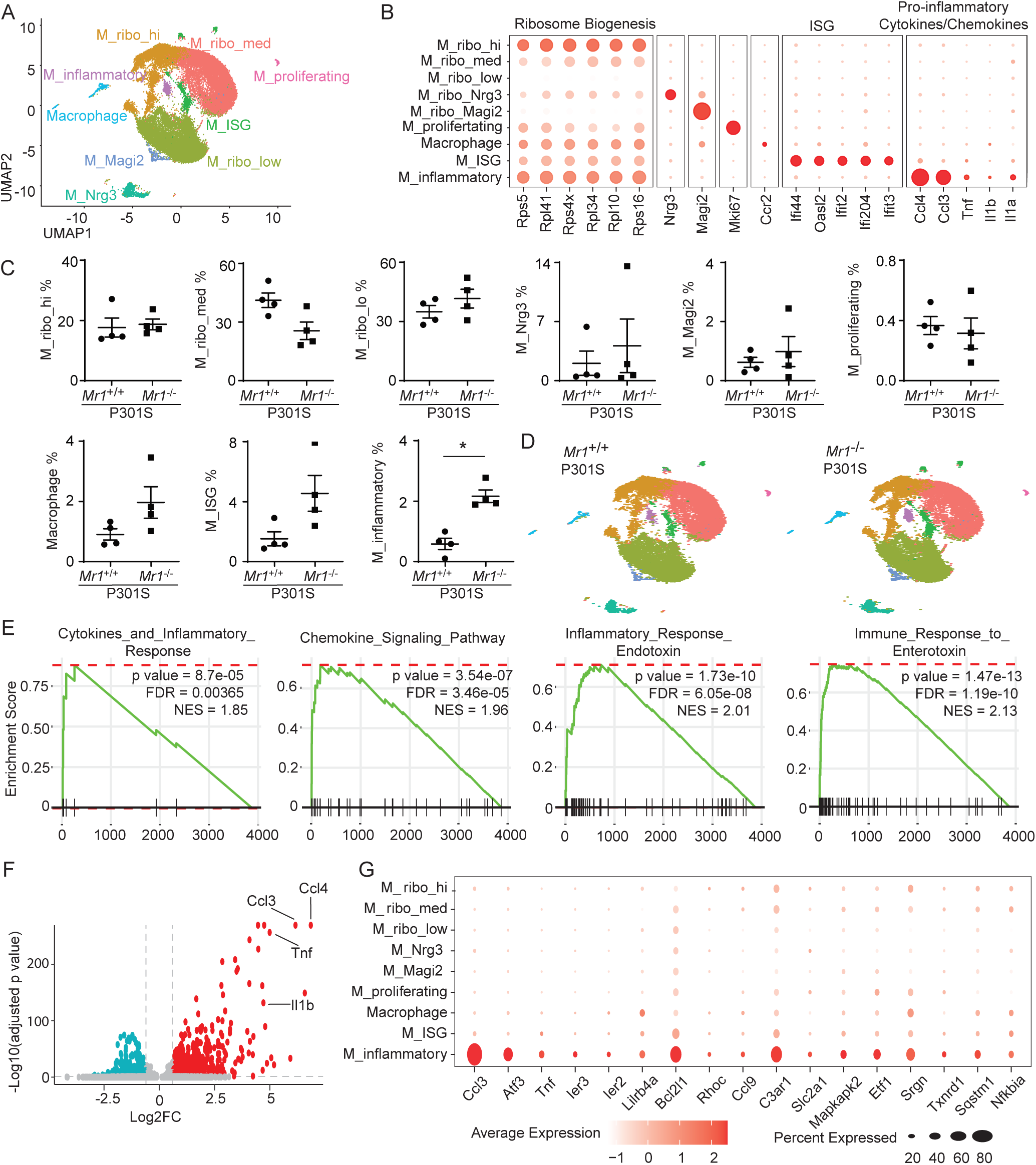
Transcriptome profiles of hippocampus microglia in 7-month-old *Mr1*^−/−^ P301S mice and control *Mr1*^+/+^ P301S mice. (**A**) UMAP plot showing different clusters of microglia/macrophage in the hippocampus of 7-month-old mice by single-nucleus RNA-seq analysis. Shown are all microglia/macrophage pooled from four 7-month-old *Mr1*^−/−^ P301S mice and four control *Mr1*^+/+^ P301S mice. (**B**) Dot plots showing expressional levels of the indicated genes of each subset. (**C**) Percentages of each microglia and macrophage subset. (**D**) UMAP plots showing different clusters of microglia/macrophage in the hippocampus of 7-month-old *Mr1*^−/−^ P301S mice and control *Mr1*^+/+^ P301S mice separately. (**E**) Enrichment of the indicated gene sets in M_inflammatory cells versus the other microglia/macrophage subsets. (**F**) Volcano plots depict genes that are significantly upregulated or downregulated in M_inflammatory cells compared to other microglia/macrophage cells. (**G**) Dot plots showing expressional levels of the indicated genes of each subset. Data are from 4 mice per group.

We performed a more thorough analysis of M_inflammatory cells. GSEA analysis revealed that these cells exhibit higher expression of genes associated with proinflammatory cytokine and chemokine pathways, as well as immune and inflammatory responses to toxic substances (Fig. 4E). Notably, the proinflammatory cytokine and chemokine genes *Tnf, Il1b, Ccl3*, and *Ccl4* were among the most significantly upregulated in M_inflammatory cells (Fig 4F). In addition to these proinflammatory mediators, these cells showed elevated expression of other genes related to responses to toxic substances (e.g. *Slc2a1, Mapkapk2, Etf1, Srgn, Txnrd1, Sqstm1, Nfkbia*), suggesting they may be exposed to higher levels of noxious substances (Fig. 4E, G). Together, M_inflammatory cells display a pro-inflammatory profile characterized by a distinct transcriptomic signature.

We also examined the transcriptomal changes of other microglia subsets in *Mr1^−/−^* P301S mice. We focused on the three major microglia subsets (M_ribo_hi, M_ribo_Med, M_ribo_low). Notable similarity was observed in the transcriptomal changes in all these three subsets between *Mr1^−/−^* P301S and control *Mr1*^+/+^ P301S mice (Fig. 5, A-F). Specifically, the genes upregulated in each microglia subset in *Mr1^−/−^*P301S mice were found to be enriched in genes related with lipoprotein assembly and clearance, ribosomal proteins, and inflammatory responses to toxic substances (Fig. 5, A-F). *Apoe*, a lipoprotein gene highly expressed in human microglia of Alzheimer’s disease patients [35], was expressed much higher in all the three main microglia subsets in *Mr1^−/−^* P301S mice compared to those in P301S mice (Fig. 5B, 5D, 5F). A few other lipoprotein assembly and clearance related genes, such as *Lpl, Apoc1, Abcg1, Nceh1, Npc2*, were also significantly upregulated in all three main microglia subsets in *Mr1^−/−^*P301S mice compared to *Mr1^+/+^* P301S mice. A variety of ribosome biogenesis genes (e.g. *Rps18, Rpl11a, Rps28, Rpl41, Rpl36a, Rpl39*) were also expressed higher in all the three main microglia subsets in *Mr1^−/−^* P301S mice, suggesting that microglia in *Mr1^−/−^* P301S mice had higher level of ribosome biogenesis and gene translation (Fig. 5, A-F). In addition, microglia in *Mr1^−/−^*P301S mice expressed higher levels of genes related to immune and inflammatory responses to toxic substances indicating that they might be exposed to higher levels of noxious substances (Fig. 5, A-F). Proinflammatory chemokines and cytokines, such as *Ccl3, Ccl4, Il1b, and Tnf*, were among the gene sets associated with responses to toxic substances. These proinflammatory chemokine and cytokine genes were upregulated in microglia of *Mr1^−/−^*P301S mice (Fig. 5B, 5D, 5F). Together, microglia in *Mr1^−/−^* P301S mice exhibit a proinflammatory profile.

**Figure 5:**
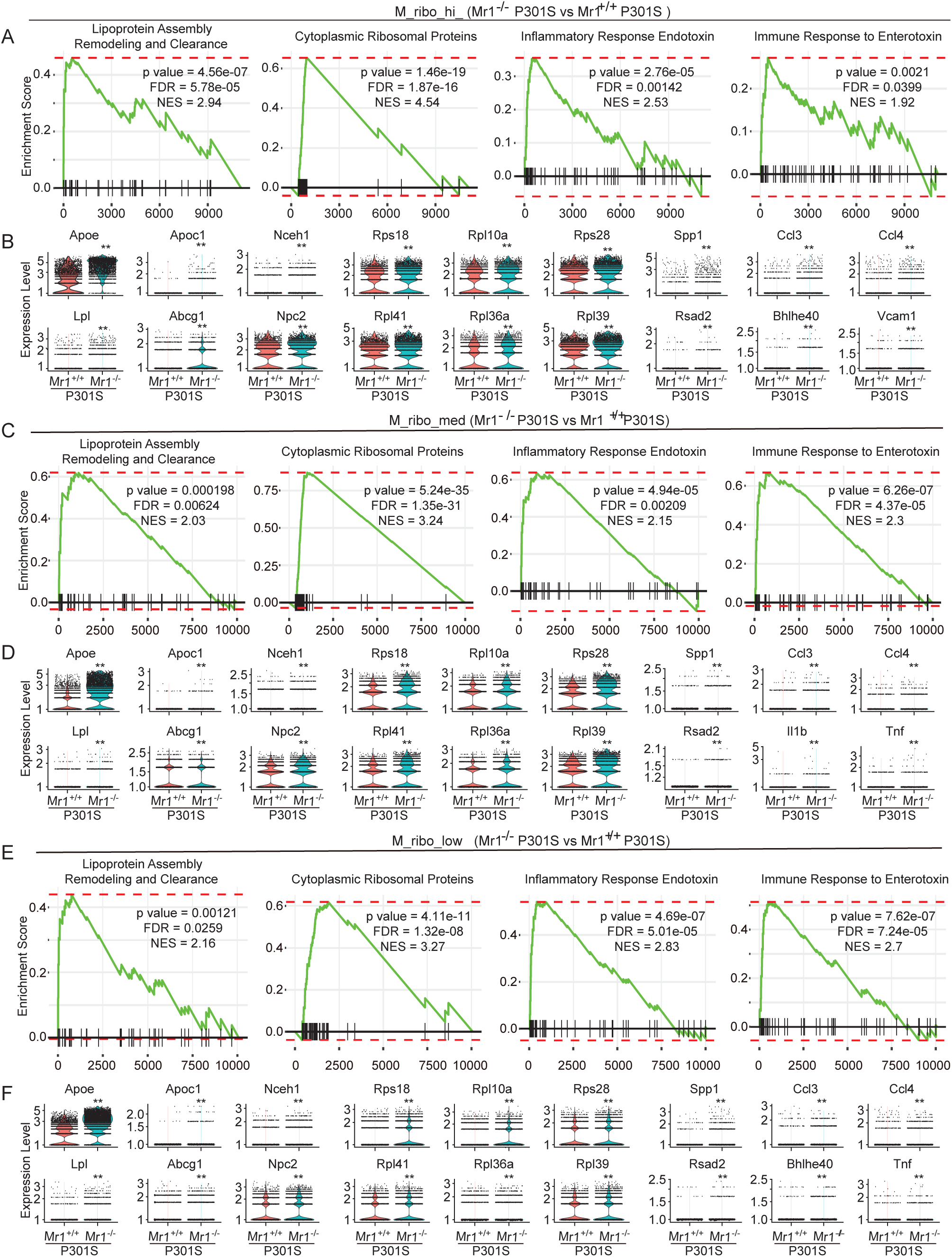
Comparison of gene expression of the main microglia subset of hippocampus microglia between 7-month-old *Mr1*^−/−^ P301S mice and control *Mr1*^+/+^ P301S mice. **(A)** Enrichment of the indicated gene sets in M_ribo_hi cells from 7-month-old *Mr1*^−/−^ P301S mice and control *Mr1*^+/+^ P301S mice. (**B**) Violin plots show expression of representative genes that were differentially expressed in M_ribo_hi cells between 7-month-old *Mr1*^−/−^ P301S mice and control *Mr1*^+/+^ P301S mice. **(C)** Enrichment of the indicated gene sets in M_ribo_med cells from 7-month-old *Mr1*^−/−^ P301S mice and control *Mr1*^+/+^ P301S mice. (**D**) Violin plots show expression of representative genes that were differentially expressed in M_ribo_med cells between 7-month-old *Mr1*^−/−^ P301S mice and control *Mr1*^+/+^ P301S mice. **(E)** Enrichment of the indicated gene sets in M_ribo_low cells from 7-month-old *Mr1*^−/−^ P301S mice and control *Mr1*^+/+^ P301S mice. (**F**) Violin plots show expression of representative genes that were differentially expressed in in M_ribo_low cells between 7-month-old *Mr1*^−/−^ P301S mice and control *Mr1*^+/+^ P301S mice. Data are from 4 mice per group.

We next performed QPCR to verify the gene expression in microglia that were FACS-sorted from the hippocampus of *Mr1^−/−^* P301S mice and control *Mr1*^+/+^P301S mice. Of note, *Apoe* was greatly upregulated in the hippocampal microglia of *Mr1^−/−^* P301S mice, compared to those in control *Mr1*^+/+^P301S mice (Fig. 6A). Expression of another lipoprotein *Apoc1* was also increased in microglia from the hippocampus of *Mr1^−/−^* P301S mice. Microglia from *Mr1^−/−^*P301S mice also exhibited increased expression of the representative ribosome biogenesis gene *Rps18*, as well as genes that were upregulated in responses to toxic substances such as *Spp1* and *Rsad1*. Transfer of MAIT cells reduced the expression of these genes in microglia from *Mr1^−/−^*P301S mice (Fig. 6B), verifying that the presence of MAIT cells repress the expression of these genes in hippocampus microglia of P301S mice. We then performed multiplex cytokine assays to verify whether the protein concentrations of proinflammatory cytokines in the brain homogenates of *Mr1^−/−^* P301S mice and control *Mr1*^+/+^P301S mice. The concentrations of proinflammatory cytokines including IL1β, IL1α, and TNF, were significantly increased in the brain homogenates of *Mr1^−/−^* P301S mice (Fig. 6C). Adoptive transfer of MAIT cells reduced the concentrations of these proinflammatory cytokines, indicating that the presence of MAIT cells repressed neuroinflammation in *Mr1^−/−^* P301S mice (Fig. 6D). Together, these data verify that MAIT cells repress microglial inflammation in P301S mice.

**Figure 6:**
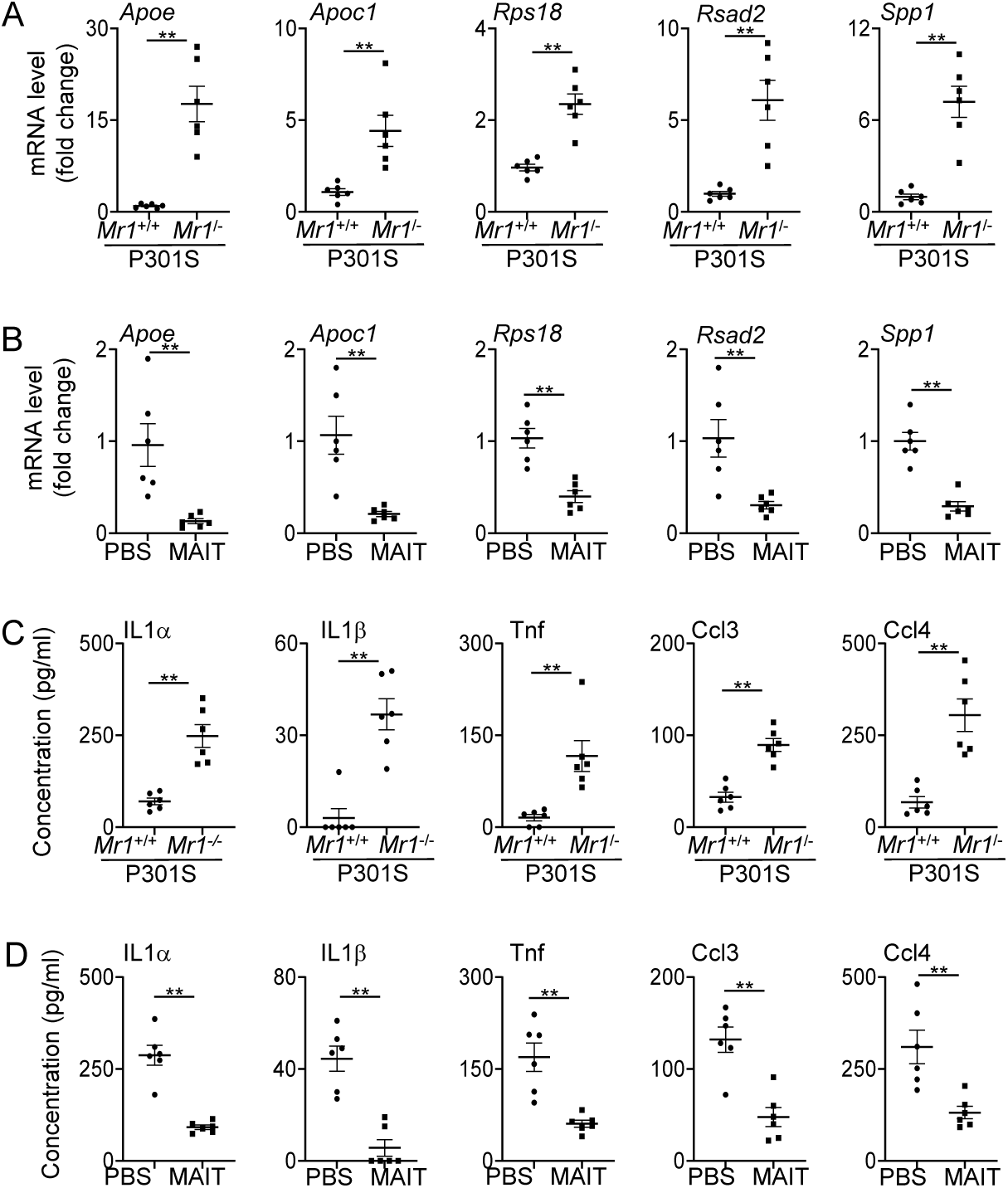
The presence of MAIT cells alters gene expression in microglia and represses proinflammatory cytokine concentrations in P301S mice. **(A)** mRNA levels of the indicated genes in microglia sorted from the hippocampus of 7-month-old *Mr1*^−/−^ P301S mice and control *Mr1*^+/+^ P301S mice. (**B**) mRNA levels of the indicated genes in microglia sorted from the hippocampus of 7-month-old *Mr1*^−/−^ P301S mice with adoptive transfer of MAIT cells or PBS. (**C**) Concentrations of the indicated cytokines in the brain homogenate of 7-month-old *Mr1*^−/−^ P301S mice and control *Mr1*^+/+^ P301S mice. (**D**) Concentrations of the indicated cytokines in the brain homogenate of 7-month-old *Mr1*^−/−^ P301S mice with adoptive transfer of MAIT cells or PBS. Data are from 6 mice per group, representative of 2 independent experiments.

## Discussion

In this study, we revealed an important role of MAIT cells in regulating tau pathology and neurodegeneration in P301S transgenic mice and investigated the underlying mechanisms. Our findings demonstrated that MAIT cells are present in the meninges and express anti-oxidant molecules. We showed that the deficiency of MAIT cells leads to meningeal barrier leakage and examined microglial dysfunction in P301S mice lacking MAIT cells. Our data together suggest an important role for MAIT cells in repressing tau related neuroinflammation and neurodegeneration.

Increasing evidence indicates that potential roles for lymphocytes particularly T cells may contribute to the regulation of brain homeostasis and neurodegeneration [5, 9–12]. MAIT cells are a unique type of innate-like T cells that are abundant in humans [36, 37]. Our previous work suggested that MAIT cells are present in the meninges of adult wildtype mice, where they express antioxidant molecules that help maintain meningeal barrier integrity [18]. However, the potential impact of meningeal barrier integrity on neurodegeneration has been poorly explored. In this study, we demonstrate the presence of MAIT cells in the meninges of P301S mice with tau pathology and confirm that these cells express antioxidant molecules to protect meningeal barrier integrity. We found that the absence of MAIT cells leads to increased microglial inflammation, exacerbating tau pathology and neurodegeneration in P301S mice. These data support the hypothesis that active maintenance of meningeal barrier integrity by meningeal-resident immune cells such as MAIT cells may be important for restricting tau pathology and neurodegeneration.

In our scRNA_seq analysis, we identified a distinct microglia subset that accumulated in the hippocampus of *Mr1*^−/−^ P301S mice, which we termed M_inflammatory. These cells exhibited proinflammatory phenotype. They mostly closely resemble the Microglia_chemokine mentioned in an earlier report [38]. However, because Microglia_chemokine cells were extremely rare in wildtype mice, detailed description of Microglia_chemokine was yet lacking in the earlier study, except that these cells express high levels of chemokines [38]. Here, we performed extensive transcriptomic analysis of M_inflammatory cells, and found that they express high levels of proinflammatory cytokines and genes related to immune and inflammatory responses to toxic substances. Our data indicate that M_inflammatory may represent a pro-inflammatory microglial subset that emerges in response to increased exposure to noxious substance. Further exploration of these cells in other pathological contexts and their specific roles in brain function and neurodegeneration might be an important avenue for future research.

Previous work indicates reactive microglia may lead to neurodegeneration in in vitro 3D human neuroimmune culture and exacerbate tau pathology in transgenic mouse models [5–9]. In our study, we observed a remarkable increase in proinflammatory cytokines in the hippocampus of *Mr1*^−/−^ P301S mice, accompanied by a pronounced proinflammatory profile of microglia. Adoptive transfer of MAIT cells reduced proinflammatory cytokine levels in the hippocampus and decreased pro-inflammatory profiles in microglia, indicating that MAIT cells repress microglial inflammation in P301S mice. Consistent with previous reports linking neuroinflammation to tau-mediated neurodegeneration [32–34], *Mr1*^−/−^ P301S mice exhibited exacerbated tau pathology and hippocampus atrophy. MAIT cell transfer reduced tau pathology and hippocampus atrophy, supporting a role of MAIT cells in repressing neurodegenerative tau pathology. [13, 15, 36, 39, 40]. The abundance of MAIT cells is highly variable and influenced by the microenvironment, particularly the microbiota [41–43]. Future research exploring the relationship between gut microbiota, MAIT cell development, and neurodegeneration could provide new insights into the mechanisms underlying neurodegenerative diseases.

### Conclusion

In conclusion, we report that MAIT cells were present in the meninges of P301 mutant human tau transgenic mice and retain expression of anti-oxidant molecules. The absence of MAIT cells exacerbates tau pathology and neurodegeneration in P301S mice. MAIT cells protect meningeal integrity and repress neuroinflammation in P301S mice. Together, these results indicate that a role of MAIT cells in regulating neurodegenerative tau pathology.

## Abbreviation

MAIT: mucosal-associated invariant T cells
MR1: MHCI-related
APC: antigen presenting cells
FACS: fluorescence activated cell sorting
snRNA-seq: single nucleus RNA sequencing
UMAP: Uniform Manifold Approximation and Projection
GSEA: Gene Set Enrichment Analysis
cDNA: Complementary DNA (cDNA)
M_inflammatory: Inflammatory Microglia
ISG: Interferon stimulated gene

## Acknowledgement

This work was supported by the U.S. National Institutes of Health Grants R01HL155021 (QY), RF1AG078459 (QY), and support to the Child Health Institute of New Jersey from the Robert Wood Johnson Foundation (Grant #74260). We thank Drs. Zhiping Pang and Arnold Rabson for help discussion.

## Authors’ contributions

Y.Z., Z.Y., X.T., J.N., G.C., and Q. Y. performed the experiments. Z.Y. performed bioinformatics analysis. Y.Z., Z.Y., and Q.Y. wrote the manuscript. All authors approved the manuscript. Data are available from the corresponding author upon request.

